# A fossilized birth-death model for fossil records lacking sampled ancestors

**DOI:** 10.64898/2026.08.13.744712

**Authors:** Jeremy M. Beaulieu, Brian C. O’Meara

## Abstract

Fossilized birth-death (FBD) models provide a powerful framework for estimating diversification from phylogenies that include fossil taxa. However, the original formulation makes a key assumption that sampled ancestors (*k*-type fossils) should be commonly observed. Beaulieu & O’Meara (2023) showed that this assumption is often violated in empirical datasets, where fossils are represented mostly or entirely as extinct terminal taxa (*m*-type fossils), which can lead to biased parameter estimates. Here, we derive the fossilized birth-death of terminal fossils (FBDT) model, an extension of the FBD, that accommodates incomplete fossil samples in which only terminal fossil occurrences are observed. We implement the model within the state-dependent speciation and extinction (SSE) framework and evaluate its performance using simulations spanning homogeneous and heterogeneous diversification scenarios. Across a wide range of fossil sampling rates, the FBDT model recovered diversification parameters that closely matched those obtained from complete fossil sample while avoiding the systematic biases that arise when sampled ancestors are unobserved. These results demonstrate that modifying the likelihood to reflect how fossil datasets are assembled provides a simple and effective extension of the FBD framework for many empirical applications.

## Introduction

Fossils are invaluable sources of information about extinct diversity and are often an integral part of phylogenetic inference for modern taxa. For instance, by explicitly modeling speciation, extinction, and fossil preservation the fossilized birth-death (FBD; Stadler, 2010) model is often cited as a way to improve the estimation of diversification rates, divergence times, and evolutionary relationships (Heath et al., 2014; Stadler et al., 2018; Louca & Pennell, 2020; Cerný et al., 2022; do Rosario Petrucci et al., 2025). Since its introduction, the FBD framework has undergone substantial development, with extensions that include accommodating stratigraphic ranges (e.g., Stadler et al., 2018), occurrence-only data (e.g., Andréoletti et al., 2022), heterogeneous diversification and fossil sampling (e.g., Mitchell et al., 2019; Beaulieu & O’Meara, 2023; Barido-Sottani & Morlon, 2026), among others (reviewed by Wright et al., 2022). These models now underlie many likelihood- and Bayesian-based analyses that integrate fossil and extant taxa into a common framework.

Although the FBD process is typically viewed as a model of diversification and fossil preservation, the model’s likelihood makes key assumptions about the fossil occurrences represented in a given dataset. Under the FBD model, fossilization events occurring on lineages represented by sampled descendants (*k*-type fossils, or sampled ancestors) and fossilization events occurring on lineages that leave no sampled descendants (*m*-type fossils, or terminal fossil occurrences) are both assumed to contribute to the observed data (Fig. 1). The likelihood interprets both the presence and absence of sampled ancestors as information about the underlying diversification and preservation processes.

**Figure 1.**
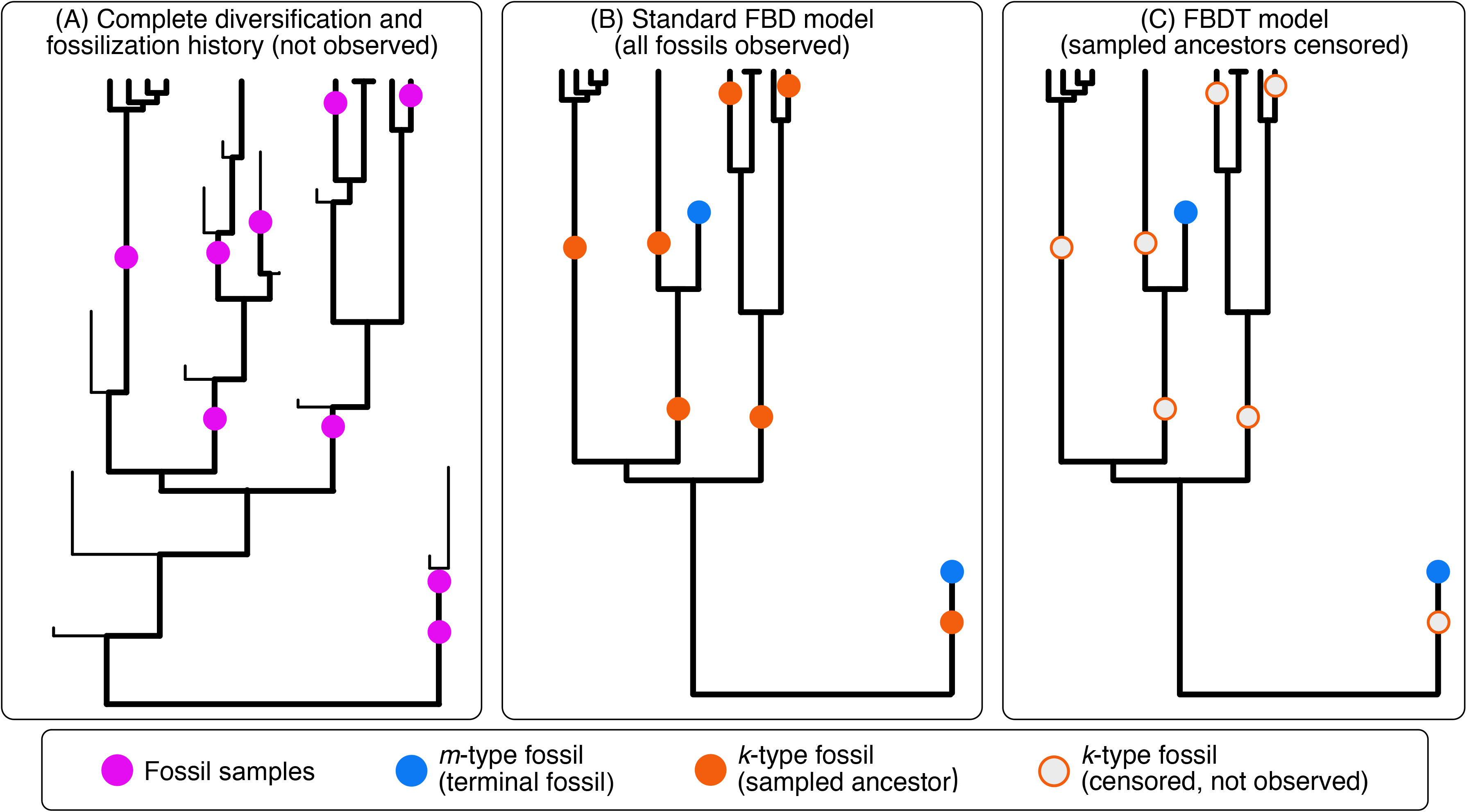
An illustration of the fossilized birth-death process and the FBDT model. (**A**) A complete diversification and fossilization sample generated under a birth-death process, including both extinct and extant lineages. This complete history is unobserved. (**B**) The sampled phylogeny obtained from (**A**), containing only sampled extant taxa and fossil occurrences. Under the standard fossilized birth-death (FBD) model, fossil samples are classified as either *m*-type fossils (blue circles), representing terminal fossil lineages with no sampled descendants, or *k*-type fossils (orange circles), representing sampled ancestors with sampled descendant lineages. (**C**) The FBDT model introduced here, in which *k*-type fossils are assumed to be censored or unobserved and only *m*-type fossils are available.

However, many empirical fossil datasets are assembled in ways that differ from the assumptions of the FBD model. Both theoretical models of fossil preservation, as well as the FBD process itself, predict that sampled ancestors should be common (Foote, 1996; Beaulieu & O’Meara 2023; Parins-Fukuchi & Saulsbury 2025). Recognizing and incorporating sampled ancestors into phylogenetic inference has long presented methodological challenges in paleontology (Fisher, 2008), and, as a result, many empirical datasets contain few or no fossil occurrences that would be interpreted as sampled ancestors. Beaulieu & O’Meara (2023) showed through simulations that applying the standard FBD likelihood to such datasets can substantially bias diversification estimates because the model interprets the biased absence of sampled ancestors as biological information rather than due to dataset assembly. Recently, do Rosario Petrucci et al. (2025) showed that FBD models remain reassuringly unbiased even with low fossil sampling rates, but they did not model the empirical bias in reducing the proportion of sampled *k*-type fossils *relative* to included *m*-type fossils. As such, their simulations do not speak to the effects of systematic absence of sampled ancestors relative to other fossils.

The handling of biased datasets is an example of a more general form of ascertainment bias, in which some kinds of observations are systematically excluded and not accounted for in the likelihood. A more familiar example lies in the analysis of discrete morphological characters. Morphological datasets have traditionally excluded invariant characters because they contain no information under parsimony and can vastly outnumber variable traits (i.e., no one scores “has bones” in a study of mammal morphological evolution). However, removing these characters creates an ascertainment bias, where analyzing only variable characters makes high rates of character evolution appear much more likely than they really are. The relevant question, therefore, is not whether models are robust to fast or slow rates of evolution, but whether they are robust to the systematic exclusion of invariant characters. Lewis (2001), building on earlier work by Felsenstein (1992), addressed this problem by deriving a likelihood that conditions on observing only variable characters.

Including morphological data can substantially improve phylogenetic inference, but only when the likelihood reflects how those data were assembled (Wright & Hillis, 2014). The issue addressed here is analogous. Incomplete fossil records that systematically exclude sampled ancestors require a likelihood whose assumptions match the available fossil data, rather than one derived for properly sampled fossil histories. While Beaulieu & O’Meara (2023) identified the statistical mismatch from biased fossil samples, unlike Lewis (2001), they did not at that time derive a likelihood appropriate for fossil datasets lacking sampled ancestors.

Here, we derive a modified fossilized birth-death model, which we refer to as the FBDT model, for datasets containing only terminal fossil occurrences. Rather than treating the absence of sampled ancestors as evidence that no such fossilization events occurred, FBDT treats fossilization events occurring on lineages with sampled descendants as unobserved or hidden. This modification leaves the underlying diversification and fossilization processes unchanged but derives a likelihood whose assumptions correspond to the available fossil data. Using simulations spanning a range of diversification and fossilization scenarios, we compare FBDT with the standard FBD likelihood and show that FBDT substantially reduces bias when analyzed datasets contain only terminal fossil occurrences.

## The FBDT model

The fossilized birth-death (FBD) model (Stadler, 2010) describes diversification as a continuous-time process in which speciation, extinction, and fossilization occur instantaneously through time. As illustrated in Figure 1, fossilization events may occur on lineages that leave no sampled descendants after the fossilization event (*m*-type fossils) or on lineages that survive beyond the fossilization event and leave sampled descendants (*k*-type fossils). Importantly, with the FBD both *m*- and *k*-type fossils are generated under the same tree-wide fossil sampling, *ψ*, such that *ψ _m_* = *ψ _k_* (Beaulieu & O’Meara, 2023). Thus, the FBD likelihood explicitly assumes that both fossil types should be observed.

As discussed by Beaulieu & O’Meara, (2023), this assumption is frequently violated in empirical datasets because sampled ancestors are often relatively or completely absent despite otherwise informative fossil records. Figure 2 illustrates the statistical consequences of applying the FBD likelihood under these conditions. When the complete fossil history is available, the likelihood surface is centered on the generating diversification parameters, regardless of the fossil sampling rate, as expected (Fig. 2A, D). However, simply removing *k*-type fossils and applying the canonical FBD likelihood shifts the likelihood surface away from the generating values (Fig. 2B, E). This behavior is not a consequence of reduced fossil sampling *per se*, but rather of applying a likelihood that assumes a variety of different fossil placements to data in which a subset is systematically missing or unobserved. As fossil sampling increases, the expected number of sampled ancestors also increases. Their absence forces the likelihood to compensate by favoring higher turnover and extinction fractions as a consequence of higher absolute extinction rates. In other words, the complete lack of *k*-type fossils is effectively telling the likelihood calculation that fossilized lineages have a very low probability of surviving long enough to become sampled ancestors.

**Figure 2.**
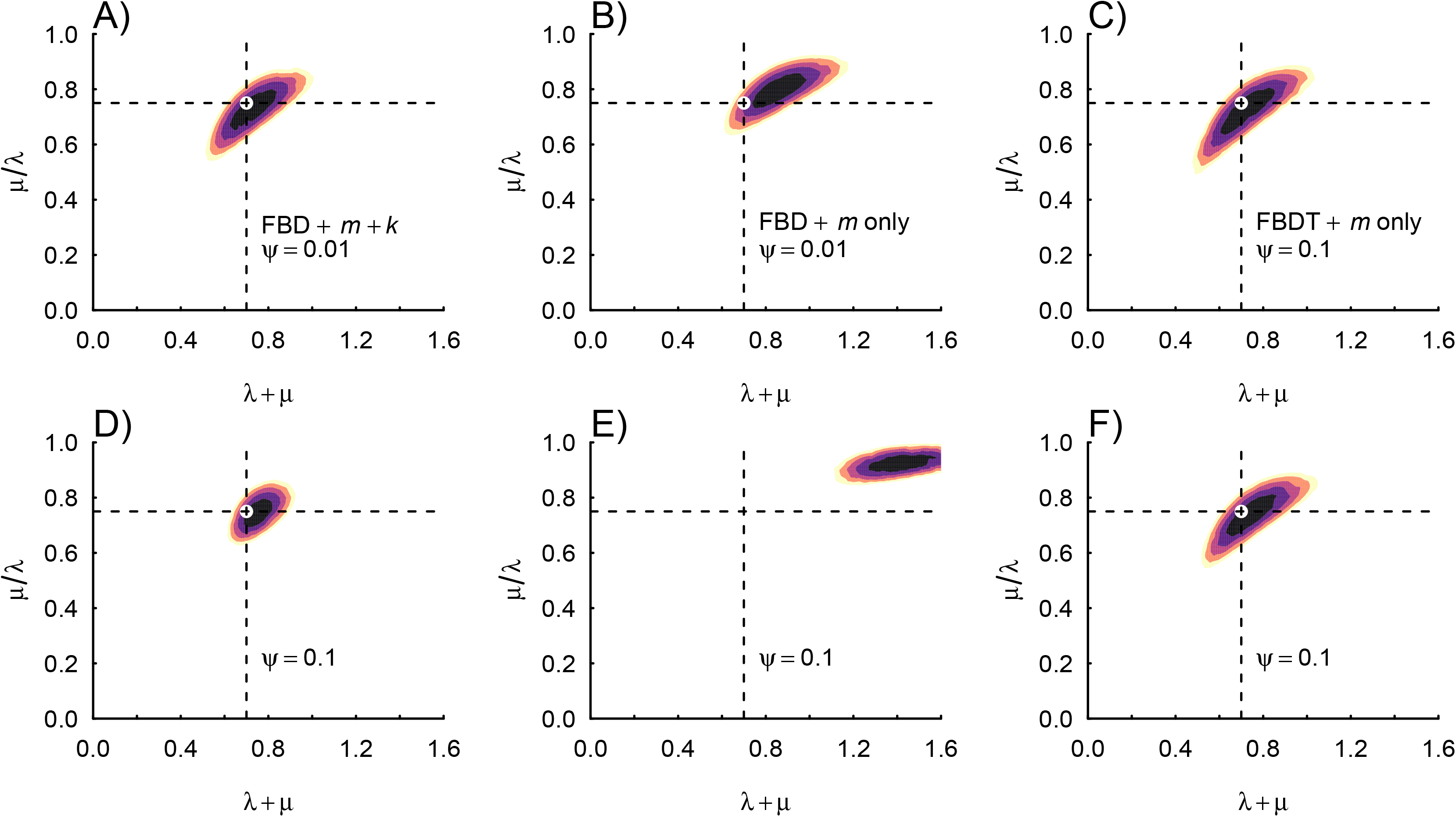
Likelihood surfaces for turnover rate (*τ*) and extinction fraction (*ε*) under alternative assumptions about fossil sampling for the same simulated tree. Each likelihood surface was evaluated by fixing *τ* and *ε* over a Latin hypercube design of 5,000 parameter combinations while optimizing the fossilization rate (*ψ*). Panels (**A**, **D**) show analyses using the complete fossil sample under the standard FBD model, (**B**, **E**) analyses of *m*-type fossils only under the canonical FBD model, and (**C**, **F**) analyses of *m*-type fossils under the FBDT model. The top row assumes *ψ* = 0.01 and the bottom row *ψ* = 0.10. Dashed vertical and horizontal lines denote the generating values of *τ* = 0.70 and *ε* = 0.75, respectively.

To provide a possible solution to this problem, we derive an extension of the canonical FBD model that explicitly accommodates incomplete fossil samples in which only *m*-type fossils are included (Fig. 1). We refer to this model hereafter as the fossilized birth-death of terminal fossils, or simply the FBDT model. The diversification process itself remains the same. Instead, following the fossilized *SSE models of Beaulieu & O’Meara (2023), we modify the branch probabilities, which includes diversification rates, state transitions, and a tree-wide fossil sampling rate, *ψ*, in the following way:

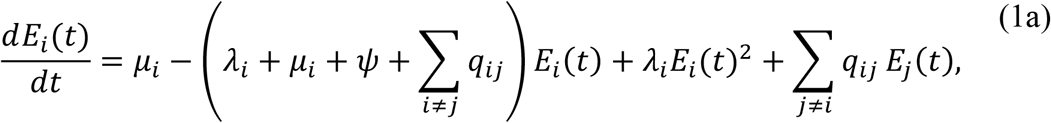

and

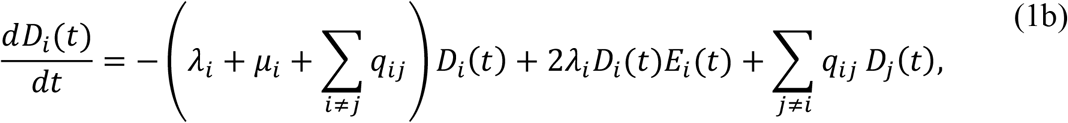

The probability *E_i_*(*t*) is the probability that a lineage starting at time *t* in state *i* leaves no sampled extinct or extant descendants at the present day (*t* = 0), and *D_i_*(*t*) is the probability of a lineage in state *i* at time *t* before the present (*t* > 0) evolved the exact observed history, including its branching structure, fossil sampling events, surviving tips, and, for models of character evolution, the observed character states.

Note that the only substantive change from the canonical FBD formulation occurs in Eq. (1b) which removes *ψ* from *D_i_*(*t*). Under the canonical FBD model, both *m*- and *k*-type fossils are possible observations and therefore contribute to the probability of the observed history. The absence of *k*-type fossils is itself informative for the canonical FBD: a history containing no *k*-type fossils requires an explanation and is evaluated against possible histories in which *k*-type fossil sampling events could have been observed. Because *k*-type fossils are not represented in the observed fossil set under the FBDT, fossilization events on lineages that have sampled descendants no longer contribute to the observed history (see Appendix A). Thus, these events are removed from the branch probability calculation whereas fossil sampling events corresponding to observed *m*-type fossils continue to contribute through *E_i_*(*t*) at the time of fossil sampling. We note that this modification does not change the underlying fossil sampling rate, *ψ*, only which fossil sampling events contribute to the probability of the observed history. Finally, we note that for character-dependent models such as HiSSE (Beaulieu & O’Meara, 2016) and MuHiSSE (Nakov et al., 2019), *i* and *j* represent combinations of observed and hidden states, whereas in MiSSE (Vasconcelos et al., 2022) they correspond only to hidden diversification regimes.

To compute the likelihood on a tree with the full model requires numerically integrating probabilities rootward using ordinary differential equations, following the same algorithm as Maddison et al. (2007) and its successors. Appendix A provides an analytical solution for the single-rate birth-death case, where we define *p*_1_^*^, to replace the *p*_1_ term of Stadler (2010) in the original FBD formulation. We demonstrate that this analytical solution produces identical log-likelihoods to the numerical ODE implementation described above (Supplementary Materials).

Before moving forward, we demonstrate the behavior of the likelihood surface when removing the fossil sampling rate term from *D_i_*(*t*). Because fossilization events along lineages with sampled descendants no longer contribute to the likelihood, the displacement of the likelihood surface observed under the canonical FBD model disappears. Incomplete fossil sampling histories nevertheless contain less information (simply due to fewer data points) than complete fossil samples, so the resulting likelihood surface is broader but remains centered on the generating diversification parameters (Fig. 2C, F). Thus, the FBDT model correctly attributes the loss of information from incomplete fossil sampling to increased uncertainty rather than to changes in the underlying diversification process.

## Recovering diversification parameters from incomplete fossil records

We evaluated the performance of the FBDT implementation in MiSSE (Vasconcelos et al., 2022) using simulations spanning both homogeneous and heterogeneous diversification scenarios across a range of fossil sampling rates (*ψ*). The MiSSE model provides a convenient test case because diversification heterogeneity is modeled through hidden states without requiring an observed character, allowing the effects of incomplete fossil sampling to be evaluated independently of trait evolution. Here, we assess whether the FBDT model recovers accurate diversification parameters from incomplete fossil samples under increasingly complex evolutionary scenarios. Throughout these simulations, sampled fossils were assumed to be placed perfectly on the generating tree so that the results reflect the statistical behavior of incomplete fossil sampling rather than uncertainty in fossil placement.

### Single-rate scenarios

We first examined the behavior of the FBDT model under a constant birth-death process across three extinction fractions (*ε* = 0.25, 0.50, and 0.75) and three fossil sampling rates (*ψ* = 0.01, 0.05, and 0.10), with a speciation rate fixed at *λ* = 0.40. Trees were simulated until they reached 200 extant taxa. For each simulated dataset, we compared parameter estimates obtained under three assumptions about the fossil record: (1) the complete, unbiased fossil sample analyzed under the canonical FBD likelihood; (2) reduced fossil sample in which a half of *k*-type fossils are removed or stratigraphic ranges were retained (fewer datapoints, more bias, though less than we see in many empirical datasets); and (3) incomplete fossil samples consisting of only *m*-type fossils analyzed using the FBDT model (least amount of data, full bias). These comparisons allowed us to assess how well FBDT recovers the results with its smallest dataset compared with canonical FBD using a large unbiased dataset and FBD using a realistically biased but medium-sized dataset.

When the complete fossil sample was used, parameter estimates for turnover (speciation rate plus extinction rate; denoted by *τ*), extinction fraction (extinction rate divided by speciation rate; denoted by *ε*), and net diversification (speciation rate minus extinction rate; denoted by *r*) under the canonical FBD likelihood were generally centered on their generating values. As expected, applying the canonical FBD likelihood directly to incomplete fossil samples produced substantial bias in the parameter estimates, particularly an upward bias in estimates of turnover and extinction fraction (Table 1). This bias generally became more pronounced as the fossil sampling rate increased, because higher sampling rates increase the expected proportion of *k*-type fossils under the canonical FBD (also see Beaulieu & O’Meara, 2023). By contrast, estimates obtained with the FBDT model remained generally close to those from the complete fossil sample across the range of generating conditions considered, indicating that the model largely corrects the systematic bias introduced when a fossil sample is incomplete. In fact the estimates from the FBDT were generally closer to the complete fossil sample than canonical FBD estimates from samples in which only a portion of the *k*-type fossils had been removed (Table 1). We also note that net diversification was comparatively robust to the completeness of the fossil sample, with smaller biases overall than those observed for turnover and extinction fraction.

**Table 1.** Summary of accuracy (bias = median of true - estimated) and precision (variance of the bias) of parameters from various simulations of the single-rate fossilized birth-death model.

| Method | $\tau$ | | $\varepsilon$ | | $r$ | | $\psi$ | |
| --- | --- | --- | --- | --- | --- | --- | --- | --- |
|  | bias | var | bias | var | bias | var | bias | var |
| $\tau = 0.50, \varepsilon = 0.25, \psi = 0.01$ | | | | | | | | |
| Full $m+k$ sample, FBD | 0.057 | 0.009 | 0.080 | 0.012 | -0.019 | 0.001 | 0.001 | <0.001 |
| 50% of $k$ fossils, FBD | 0.091 | 0.010 | 0.123 | 0.012 | -0.031 | 0.001 | -0.004 | <0.001 |
| $m$-only sample, FBD | 0.127 | 0.011 | 0.173 | 0.012 | -0.041 | 0.001 | -0.008 | <0.001 |
| Strati. ranges, FBDR | 0.026 | 0.009 | 0.046 | 0.012 | -0.016 | 0.001 | 0.001 | <0.001 |
| $m$-only sample, FBDT | 0.042 | 0.016 | 0.067 | 0.025 | -0.019 | 0.002 | 0.003 | 0.021 |
| $\psi = 0.05$ | | | | | | | | |
| Full $m+k$ sample, FBD | 0.015 | 0.007 | 0.009 | 0.009 | 0.000 | 0.001 | -0.001 | <0.001 |
| 50% of $k$ fossils, FBD | 0.088 | 0.010 | 0.105 | 0.010 | -0.018 | 0.001 | -0.024 | <0.001 |
| $m$-only sample, FBD | 0.222 | 0.015 | 0.237 | 0.010 | -0.053 | 0.001 | -0.045 | <0.001 |
| Strati. ranges, FBDR | -0.030 | 0.005 | -0.051 | 0.007 | 0.010 | 0.001 | 0.002 | <0.001 |
| $m$-only sample, FBDT | 0.021 | 0.016 | 0.030 | 0.024 | -0.002 | 0.002 | -0.003 | 0.080 |
| $\psi = 0.10$ | | | | | | | | |
| Full $m+k$ sample, FBD | 0.003 | 0.004 | -0.009 | 0.007 | -0.001 | 0.001 | 0.000 | <0.001 |
| 50% of $k$ fossils, FBD | 0.088 | 0.008 | 0.098 | 0.009 | -0.025 | 0.001 | -0.049 | <0.001 |
| $m$-only sample, FBD | 0.310 | 0.015 | 0.310 | 0.009 | -0.071 | 0.001 | -0.092 | <0.001 |
| Strati. ranges, FBDR | -0.044 | 0.003 | -0.070 | 0.005 | 0.009 | 0.001 | 0.010 | <0.001 |
| $m$-only sample, FBDT | 0.043 | 0.017 | 0.044 | 0.025 | -0.013 | 0.002 | -0.016 | 0.056 |
| $\tau = 0.60, \varepsilon = 0.50, \psi = 0.01$ | | | | | | | | |
| Full $m+k$ sample, FBD | 0.007 | 0.011 | 0.029 | 0.013 | -0.006 | 0.001 | <0.001 | <0.001 |
| 50% of $k$ fossils, FBD | 0.046 | 0.011 | 0.063 | 0.012 | -0.015 | 0.001 | -0.004 | <0.001 |
| $m$-only sample, FBD | 0.088 | 0.012 | 0.096 | 0.011 | -0.025 | 0.001 | -0.007 | <0.001 |
| Strati. ranges, FBDR | -0.022 | 0.012 | -0.020 | 0.016 | 0.003 | 0.001 | <0.001 | <0.001 |
| $m$-only sample, FBDT | 0.010 | 0.017 | 0.024 | 0.022 | -0.008 | 0.002 | -0.001 | 0.001 |
| $\psi = 0.05$ | | | | | | | | |
| Full $m+k$ sample, FBD | 0.036 | 0.008 | 0.028 | 0.008 | -0.005 | 0.001 | -0.002 | <0.001 |
| 50% of $k$ fossils, FBD | 0.139 | 0.010 | 0.132 | 0.006 | -0.030 | 0.001 | -0.023 | <0.001 |
| $m$-only sample, FBD | 0.329 | 0.014 | 0.238 | 0.005 | -0.059 | 0.001 | -0.040 | <0.001 |
| Strati. ranges, FBDR | -0.048 | 0.007 | -0.061 | 0.009 | 0.012 | 0.001 | 0.004 | <0.001 |
| $m$-only sample, FBDT | 0.075 | 0.019 | 0.063 | 0.019 | -0.014 | 0.001 | -0.011 | 0.012 |
| $\psi = 0.10$ | | | | | | | | |
| Full $m+k$ sample, FBD | 0.003 | 0.005 | 0.013 | 0.005 | -0.006 | 0.001 | -0.001 | <0.001 |
| 50% of $k$ fossils, FBD | 0.137 | 0.008 | 0.138 | 0.004 | -0.034 | 0.001 | -0.047 | <0.001 |
| $m$-only sample, FBD | 0.436 | 0.014 | 0.290 | 0.003 | -0.080 | 0.001 | -0.084 | <0.001 |
| Strati. ranges, FBDR | -0.089 | 0.003 | -0.083 | 0.004 | 0.009 | 0.001 | 0.015 | <0.001 |
| $m$-only sample, FBDT | 0.026 | 0.016 | 0.039 | 0.018 | -0.012 | 0.001 | -0.013 | 0.024 |
| $\tau = 0.70, \varepsilon = 0.75, \psi = 0.01$ | | | | | | | | |
| Full $m+k$ sample, FBD | 0.036 | 0.010 | 0.031 | 0.006 | -0.005 | 0.001 | -0.001 | <0.001 |
| 50% of $k$ fossils, FBD | 0.101 | 0.010 | 0.054 | 0.005 | -0.009 | 0.001 | -0.004 | <0.001 |
| $m$-only sample, FBD | 0.159 | 0.010 | 0.077 | 0.005 | -0.017 | 0.001 | -0.006 | <0.001 |
| Strati. ranges, FBDR | -0.008 | 0.011 | -0.020 | 0.008 | 0.005 | 0.001 | <0.001 | <0.001 |
| $m$-only sample, FBDT | 0.059 | 0.010 | 0.027 | 0.006 | -0.005 | 0.001 | -0.002 | <0.001 |
| $\psi = 0.05$ | | | | | | | | |
| Full $m+k$ sample, FBD | 0.036 | 0.006 | 0.020 | 0.003 | -0.006 | <0.001 | -0.002 | <0.001 |
| 50% of $k$ fossils, FBD | 0.191 | 0.007 | 0.090 | 0.002 | -0.023 | <0.001 | -0.021 | <0.001 |
| $m$-only sample, FBD | 0.432 | 0.012 | 0.156 | 0.001 | -0.043 | <0.001 | -0.036 | <0.001 |
| Strati. ranges, FBDR | -0.141 | 0.004 | -0.086 | 0.003 | 0.016 | <0.001 | 0.012 | <0.001 |
| $m$-only sample, FBDT | 0.061 | 0.011 | 0.043 | 0.005 | -0.008 | 0.001 | -0.008 | <0.001 |
| $\psi = 0.10$ | | | | | | | | |
| Full $m+k$ sample, FBD | 0.031 | 0.005 | 0.015 | 0.004 | -0.004 | 0.001 | -0.001 | <0.001 |
| 50% of $k$ fossils, FBD | 0.247 | 0.008 | 0.108 | 0.002 | -0.027 | <0.001 | -0.046 | <0.001 |
| $m$-only sample, FBD | 0.719 | 0.020 | 0.191 | 0.001 | -0.056 | <0.001 | -0.079 | <0.001 |
| Strati. ranges, FBDR | -0.140 | 0.003 | -0.095 | 0.004 | 0.019 | 0.001 | 0.027 | <0.001 |
| $m$-only sample, FBDT | 0.079 | 0.013 | 0.045 | 0.006 | -0.011 | 0.001 | -0.018 | 0.001 |

Increasing the fossil sampling rate generally reduced the variance of parameter estimates for both the canonical FBD and FBDT analyses, with the greatest improvements occurring when extinction fractions and fossil sampling rates were moderate to high (Table 1). However, when the extinction rate was low relative to the speciation rate (*ε* = 0.25), parameter estimates exhibited increased variance overall, despite low bias, even under the highest fossil sampling rates. Low extinction rates produce relatively fewer extinct lineages and, consequently, fewer fossil occurrences that contribute information about the diversification process. Indeed, as extinction fraction increased, the variance of the FBDT estimates approached that obtained from the complete fossil sample, reflecting the greater proportion of fossil occurrences represented by *m*-type fossils. This pattern was particularly pronounced for estimates of the fossil sampling rate itself, *ψ*, which were considerably more variable under FBDT at low extinction fractions.

Finally, as first reported by Beaulieu & O’Meara (2023), reducing fossil occurrence data to stratigraphic ranges generally produced a downward bias in estimates of turnover and extinction fraction, particularly as fossil sampling increased, relative to both a complete fossil sample and the FBDT analyses. This showed that collapsing fossil occurrences into ranges results in some informational loss contained in the full pattern of fossil sampling. Specifically, the model does not allow unobserved speciation or extinction events to occur along a defined stratigraphic range (Beaulieu & O’Meara, 2023). Note that these are concrete shifts in the mean, not merely an increase in variance.

### Heterogeneous diversification

We next evaluated the FBDT model under three heterogeneous diversification scenarios representing progressively more challenging inference problems: differences in turnover rate alone (Scenario 1; Fig 3A-C), simultaneous differences in turnover rate and extinction fraction while maintaining nearly identical net diversification (Scenario 2; Fig 3D-F), and a homogeneous diversification process despite fitting mostly heterogeneous models (Scenario 3; Fig 3G-I). As in the single-rate simulations, each scenario was analyzed using (1) the complete fossil sample under the canonical FBD likelihood, (2) reduced fossil samples in which some or all *k*-type fossils were omitted or fossil occurrences were represented as stratigraphic ranges, and (3) incomplete fossil samples analyzed using FBDT.

**Figure 3.**
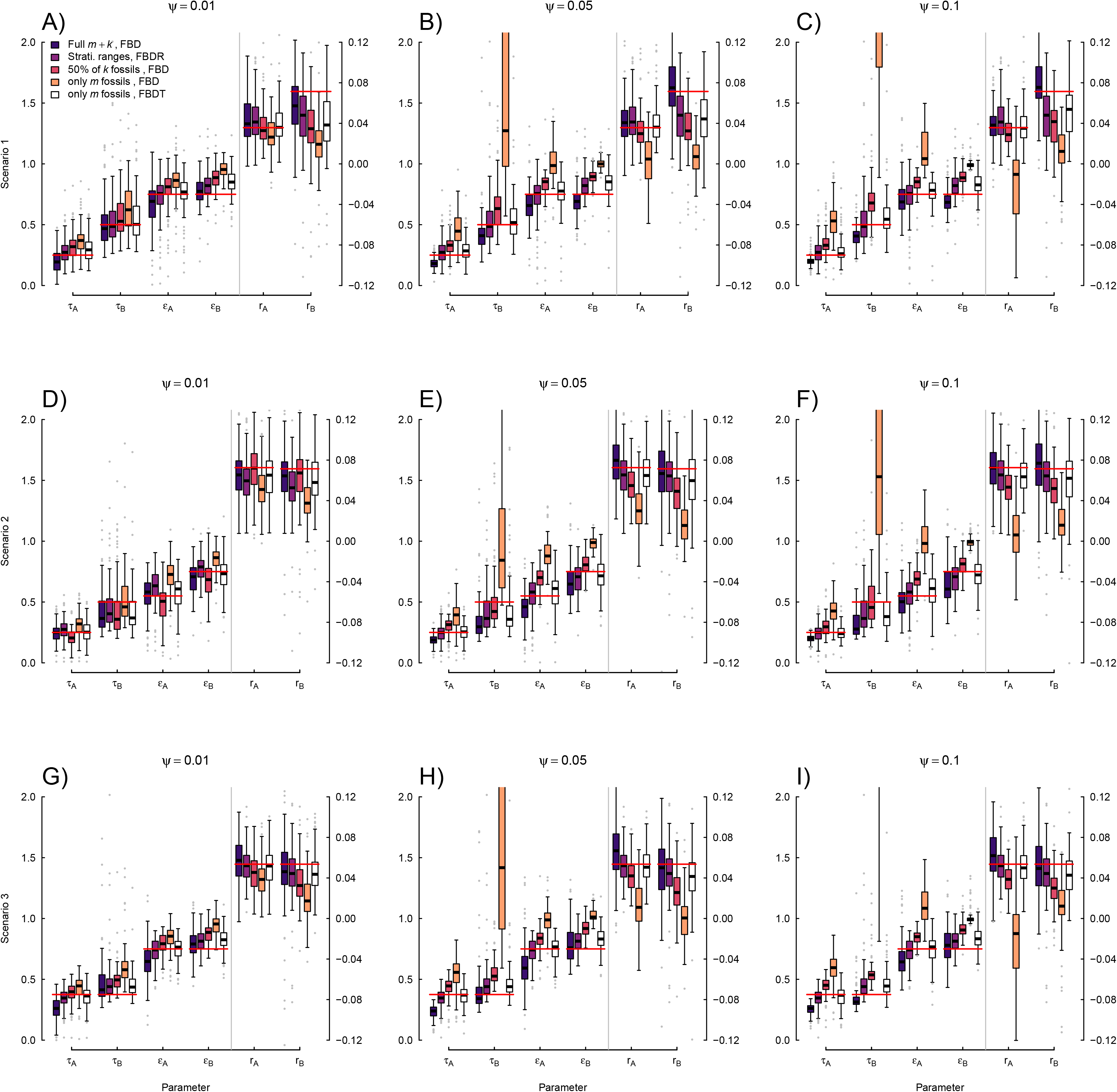
Distribution of model-averaged parameter estimates under three heterogeneous diversification scenarios. Panels **A-C** show simulations in which regime B had twice the turnover rate of regime A (*τ_A_*=0.25 events/Myr and *τ_B_* =0.50 events/Myr), while both regimes shared an extinction fraction of 0.75. Panels **D-F** show simulations in which regime B again had twice the turnover rate of regime A, but regime A had a lower extinction fraction (*ε_A_*=0.55 and *ε_B_*=0.75), which set the net diversification rate, *r*, to be nearly identical among the two regimes. Panels **G-I** show simulations under a homogeneous generating model with identical turnover rates (*τ_A_*=*τ_B_*=0.375 events/Myr) and extinction fractions (*ε_A_*=*ε_B_*=0.75). Columns correspond to increasing fossilization rates (*ψ* = 0.01, 0.05, and 0.10). Each dataset was analyzed under alternative assumptions about fossil sampling, including the proposed FBDT model, which assumes only *m*-type fossils are observed. Estimates were model-averaged across four models allowing turnover rate and extinction fraction to vary. Red horizontal lines indicate the generating values of each parameter; the secondary *y*-axis gives the corresponding scale for net diversification. The model and combinations used were the original FBD with all fossils, FBD with stratigraphic sampling (FBDR) with the ranges derived from all fossils, original FBD with only 50% of *k*-fossils included, original FBD with zero *k*-fossils included, and FBDT with zero *k-*fossils included.

For each simulation replicate, within each scenario, we fit four MiSSE models that broadly capture the complexity of the scenarios. Specifically, we fit a single regime model, a model that only allows turnover rate to vary (i.e., *τ_A_* ≠ *τ_B_*), a model that only allowed extinction fraction to vary (i.e., *ε_A_* ≠ *ε_B_*), and a model that allowed both to vary (i.e., *τ_A_* ≠ *τ_B_* and *ε_A_* ≠ *ε_B_*). Each model was evaluated using a two-step optimization routine. The first step consists of a bounded stochastic simulated annealing run for 5000 iterations, followed by a bounded subplex routine that searches parameter space until the maximum likelihood is found. Model-averaged parameter estimates were then calculated using AIC weights after removing effectively redundant models from the candidate set (see discussion in Vasconcelos et al., 2022). Model-averaging is particularly advantageous in this context because parameter estimates are not based on a single best-supported model, allowing support for multiple diversification scenarios to be reflected in the final estimates (Burnham & Anderson, 2002).

As in the single-rate simulations, we first compared analyses based on unbiased fossil samples for both the original FBD and the FBD with stratigraphic range only (FBDR: Stadler et al., 2018) with those based on incomplete fossil records. Overall, reanalyzing incomplete fossil samples with the canonical FBD likelihood again resulted in turnover and extinction fraction being systematically biased (Fig. 3). By contrast, parameter estimates obtained with the FBDT model closely tracked those from the complete fossil sample while exhibiting the modest increase in uncertainty expected from the loss of information from a reduced fossil set. As with the single-rate case, increasing fossil sampling primarily reduced the variance of parameter estimates rather than substantially changing their mean values.

Scenario 1, in which hidden states differed only in turnover rate (i.e., *τ_A_*=0.25 events/Myr and *τ_B_* =0.50 events/Myr), was recovered with relatively little bias under the FBDT model and increasing fossil sampling reduced uncertainty around estimates of both turnover regimes. Scenario 2 proved to be particularly more difficult regardless of how the fossil record was represented. Although the FBDT model consistently outperformed analyses that applied the canonical FBD likelihood to incomplete fossil samples, estimates associated with the higher-turnover regime remained biased downward. Because turnover and extinction fraction changed together (i.e., *τ_A_*=0.25 events/Myr and *τ_B_*=0.50 events/Myr; *ε_A_*=0.55 and *ε_B_*=0.75), while net diversification remained nearly constant, this scenario represents an inherently difficult inference problem in which multiple diversification processes produce similar branching patterns.

Finally, scenario 3 evaluated whether the FBDT model would spuriously infer diversification heterogeneity when the generating process was actually homogeneous (i.e., *τ_A_*=*τ_B_*=0.375 events/Myr; *ε_A_*=*ε_B_*=0.75). This is analogous to a Type I error in classical hypothesis testing, in which a more complex model is favored despite the absence of true rate heterogeneity. Despite fitting models capable of accommodating multiple hidden diversification regimes, parameter estimates for the two hidden states generally remained similar, indicating little tendency to infer false diversification shifts simply because additional model complexity was available. Although useful as a diagnostic of model behavior, we emphasize that this scenario is unlikely to represent most empirical systems. Diversification across lineages is almost certainly heterogeneous to some extent, although the underlying sources of that heterogeneity are unlikely to be fully captured by any particular diversification model.

On the whole, these simulations demonstrate that the main advantage of the FBDT model is not that it eliminates all sources of estimation error. Instead, it largely recovers the parameter estimates obtained from complete fossil histories while allowing for datasets with biased fossil records. Across both homogeneous and heterogeneous diversification scenarios, estimates obtained with FBDT consistently remained much closer to those from the complete fossil history than did analyses that ignored missing fossil occurrences or reduced fossil occurrence data to stratigraphic ranges. The remaining uncertainty reflected the amount of information available in the fossil record and the inherent difficulty of the underlying diversification scenario.

## Discussion

Fossils provide one of the few direct windows into life’s history. One of us (BCO) was recently reminded of this while admiring the spectacular ammonite fossils exposed along the Jurassic Coast at Lyme Regis. Yet the very incompleteness that makes the fossil record scientifically fascinating also makes it a challenge *statistically*. The canonical FBD model, like all variants of the FBD model family, necessarily simplifies the fossilization process by assuming homogeneous fossil sampling through time and among lineages. We know this assumption is violated in many groups. For example, much of the insect fossil record is concentrated in discrete amber deposits that are unevenly distributed through time and space (Poinar, 1993). There is also active debate about whether apparent extinction events often center on whether observed gaps reflect biological turnover or simply intervals lacking fossil preservation (Holland et al., 2025).

The FBDT model shares these simplifying assumptions with the canonical FBD model. Its contribution, therefore, is not to provide a universally more realistic model of fossilization, but, rather, to address one specific and common source of model misspecification: empirical datasets in which sampled ancestors are systematically under sampled or absent completely. The simulations presented here demonstrate that this seemingly modest modification has important statistical consequences. The FBDT largely recovers the parameter estimates obtained from complete fossil histories while correctly representing the additional uncertainty associated with incomplete fossil samples. It does not recover information that has been lost through incomplete fossil sampling – it prevents that loss of information from being misinterpreted as evidence for different diversification dynamics.

We hasten to point out that the FBDT should not be viewed as replacing the canonical FBD model. When sampled ancestors can be reliably identified and included, the canonical formulation remains the appropriate model. Similarly, the fossilized birth-death range (FBDR) model (Stadler et al., 2018) remains the natural choice when fossil occurrences are represented by first and last appearances. These models address different representations of the fossil record. The FBDT developed here accommodates datasets in which only terminal fossil occurrences are represented. Choosing among these models should, therefore, depend on how the fossil data have been assembled rather than on any notion that one model is universally preferable.

Of course, in practice, the natural question arises of what to do when *k*-type fossils are undersampled but not completely absent. Across a wide range of simulation conditions under the canonical FBD, *k*-type fossils are expected to make up a substantial fraction of all fossil occurrences, often 50% or more (see Fig. 3 in Beaulieu & O’Meara, 2023; Zhang et al., 2016). However, many empirical datasets contain a much smaller proportion, reflecting a substantial bias in what fossils are included. Neither the FBD nor the FBDT model is ideally suited in this situation. On the one hand, the canonical FBD sees the biased sampling of *k*-type fossils as data, and, on the other, the FBDT requires removing the small number of *k*-type fossils that are present.

One simple solution would be to discard the small sample of *k*-fossils, but to us throwing out data justifiably feels wrong. Other solutions might involve further model developments in which distinct separate fossil preservation rates for *k*- and *m*-type fossils are allowed (i.e., *ψ _m_* and *ψ _k_* not required to be equal), thereby allowing all observed fossils to contribute in some way to the likelihood. Whether such a model is possible, and/or would be formally identifiable without constraints, is not obvious and seems analogous to jointly estimating speciation, extinction, and sampling fraction, which is unidentifiable (Stadler, 2013). Even if separate sampling rates are formally identifiable, they may be difficult to estimate in practice when only a handful of *k*-type fossils are available to inform their sampling rate. Until such a model is developed, analyses containing a mixture of fossil types require choosing between the FBD and FBDT likelihoods. Given that *k*-type fossils are expected to constitute a substantial fraction of fossil occurrences under the FBD process, a useful rule of thumb may be to use the canonical FBD when *k*-type fossils comprise approximately half or more of the observed fossils, and FBDT when they are less than that. To do so, one would have to delete the few observed *k*-type fossils that are available. Based on the simulations (Table 1), more accurate estimates are obtained using FBDT with *m*-type fossils only than from FBD with *m*-type and an undersampling of *k*-type fossils, even though the latter has more data. We do emphasize, however, that this choice cannot be made using AIC, Bayes factors, or similar model-selection criteria because the two likelihoods would be applied to different representations of the underlying data.

We implemented the FBDT likelihood within the *hisse* package, allowing it to be applied to MiSSE (Vasconcelos et al., 2022), BiSSE (Maddison et al., 2007), HiSSE (Beaulieu & O’Meara, 2016), and other related state-dependent diversification models (e.g., MuHiSSE, Nakov et al., 2019). As with the existing FBD implementations, these analyses operate on a fixed phylogeny rather than jointly estimating topology and diversification parameters. Some have argued that for any question involving a phylogeny the topology, branch lengths, divergence times, and any parameters of interest, such as diversification rates, should inferred jointly (Tribble et al., 2026). Although the FBDT likelihood could, in principle, be incorporated into joint tree-inference frameworks such as *RevBayes* (Höhna et al., 2016), we view these problems as largely separate. Phylogenetic inference is itself a computationally challenging problem, and jointly estimating trees together with diversification processes substantially increases both computational burden and model complexity. In fact, many comparative analyses continue to estimate diversification parameters on a fixed tree, an approach that FBDT readily accommodates.

Some may wonder why a new model is necessary when Bayesian priors could instead prohibit sampled ancestors (e.g., Matzke & Wright, 2016). In our view, this addresses a different statistical problem. Priors restrict the parameter space but do not modify the likelihood to reflect the observation process that generated the data. To use a simple analogy, one could model the outcome of a fair coin by assuming a fair six-sided die while imposing a prior that only allows outcomes of rolling a one or two. Although this prior construction could, in principle, recover the correct outcomes, it is a far less natural statistical description than simply using a Bernoulli or binomial model. Similarly, stratigraphic ranges could, in principle, be accommodated through increasingly restrictive priors on fossil occurrences, yet the FBDR model instead derives a likelihood that directly reflects how those data are represented. The FBDT model follows the same philosophy. Rather than relying on priors to compensate for the absence of sampled ancestors, it conditions the likelihood on the type of fossil data that are actually available. This distinction is particularly important in diversification analyses, where the available information is often limited to branching times, fossil occurrences, and occasionally one or a few observed characters. Under these conditions, prior assumptions can exert substantial influence on posterior estimates, making it especially desirable for the likelihood itself to accurately describe the observation process.

Several limitations remain. First and foremost is that the FBDT model assumes a homogeneous fossilization process across the entire tree. In practice, fossilization rates are expected to vary among clades, through time, across environments, and among geographic regions (e.g., Wagner & Marcot, 2013; Holland, 2016; Barido-Sottani et al., 2019; Andréoletti et al., 2026). Extending the FBD family of models to accommodate clade-specific, time-varying, or jointly varying fossilization processes represents a natural direction for future work (see Barido-Sottani & Morlon, 2026). More generally, as with all diversification models, parameter identifiability and precision continue to present important challenges. Although diversification is often parameterized in terms of speciation rate, extinction rate, turnover, extinction fraction, or net diversification, only two of these quantities are independently estimable. Consequently, some parameters, particularly extinction rate, may exhibit broad likelihood surfaces even when equivalent parameterizations yield identical likelihoods and narrower surfaces for the new parameters. Uncertainty in parameter estimates warrants caution and a thorough examination of the precision of the estimates or other ways to examine the likelihood surface (e.g., Boyko & O’Meara, 2024), or through frameworks naturally fit for providing estimates of uncertainty like Bayesian approaches. Finally, diversification models are subject to well-known issues of identifiability (e.g., Kubo & Iwasa, 1995; Louca & Pennell, 2020), in addition to other systematic biases (e.g., Beaulieu & O’Meara, 2026), users should carefully evaluate uncertainty through likelihood profiling, Bayesian posterior distributions, or other approaches.

Taken together, the FBD family of models now accommodates several common representations of the fossil record (reviewed by Wright et al., 2022). The canonical FBD model assumes an unbiased observation of fossil occurrences, including sampled ancestors (Stadler et al. 2010). The occurrence birth-death process (OBDP) accommodates phylogenetically unassignable fossil occurrences (Andréoletti et al., 2022). The FBDR model accommodates fossil records summarized as stratigraphic ranges (Stadler et al., 2018). The FBDT model introduced here accommodates datasets in which only terminal fossil occurrences are represented. We expect this family of models to continue expanding as new statistical formulations are developed to better match the diverse ways in which empirical fossil datasets are being assembled.

## Appendix A: Analytical solution for the single-rate FBDT likelihood

The ordinary differential equations presented in the main text allow for diversification rates to vary among combinations of observed or hidden states. Here we consider the special case of a single-rate fossilized birth-death process. We begin by dropping the state transitions from the general formulation in Eq. (1) in the main text, which gives:

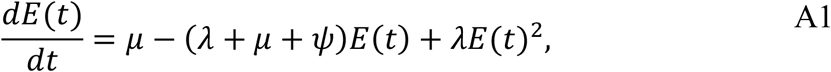

and

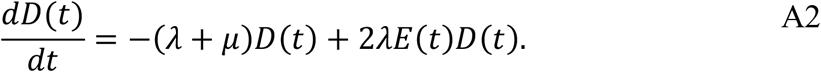

The parameters *λ*, *μ*, and *ψ* denote speciation, extinction, and the tree-wide fossil sampling rate, respectively. As in the canonical FBD model, *E*(*t*) is the probability that a lineage beginning at time *t* leaves no sampled extinct or extant descendants, whereas *D*(*t*) is the probability that a lineage beginning at time *t* has one sampled extant descendant. Because Equation A1 is identical to the corresponding “master” equation of Stadler (2010), and does not depend on *D*(*t*), its analytical solution is unchanged. We therefore retain the original expression for *p*_0_(*t*) [see Eq. (1) in Stadler 2010, pg. 398].

The expression *p*_1_(*t*) of Stadler [2010, Eq. (2), pg. 398] is derived from the following “master” equation:

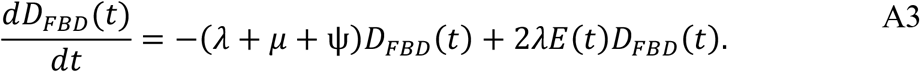

Because the only difference between Equation A2 and Equation A3 is the term −*ψD_FBD_*(*t*), we can transform *D_FBD_*(*t*) to remove it. We note that removing this term does not assume that *k*-type fossils do not occur – these events occur at a rate represented by *ψ*. But they are unobserved and so are not data that is used (in the same way that the taxa undoubtedly have traits, but for models that do not model character evolution trait data play no role). If we define *G*(*t*) = *f*(*t*)*D_FBD_*(*t*), differentiation using the product rule introduces the additional term *f*′(*t*)*D_FBD_*(*t*)such that,

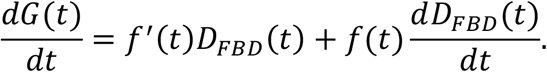

Substituting Equation A3 gives:

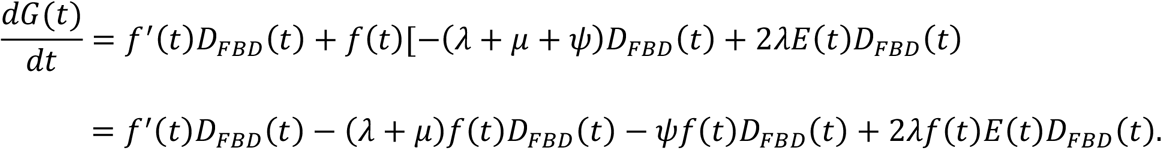

Note that the additional term, *f*′(*t*)*D_FBD_*(*t*), introduced by differentiating will cancel the −*ψf*(*t*)*D_FBD_*(*t*) term when *f*′(*t*) = *ψf*(*t*). This is satisfied by setting *f*(*t*) = *e^ψt^* since

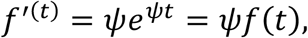

and, so, we define

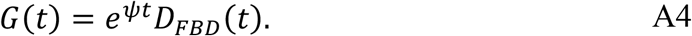

To verify, by setting *f*(*t*) = *e^ψt^* into the expression for 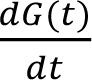 the terms involving *ψ* cancel out, leaving

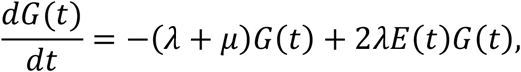

which is exactly Equation A2 describing our FBDT branch probability. Thus, we can modify *p*_1_(*t*) of Stadler (2010) as

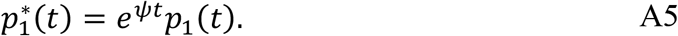

We also verified that, when *k* = 0, substituting *p*^*^_1_ into Eq. (5) of Stadler (2010; pg. 400) produces identical log-likelihoods to the ODE implementation described in the main text (Supplementary Materials).

## Data availability

All scripts and substantial outputs are available at https://github.com/thej022214/Fossils_Kcensor_BO/releases/tag/2026viii13

## Author contributions

J.M.B. developed and implemented the new model; both contributed to posing the problem, writing the manuscript, testing the method, and figures.

## Acknowledgments

We thank Andy Alverson, Diego Paredes-Burneo, and other members of the Beaulieu lab for edits that have improved the manuscript, as well as Emmett O’Meara for comments on the Appendix. We would also like to thank David Bapst for spurring fruitful discussion regarding sampled ancestors. JMB would also like to acknowledge that discussions and arguments with Tomo Parins-Fukuchi, James Pease, James Saulsbury, and Nathanael Walker-Hale regarding diversification rates, age-rate scaling, and fossils reignited his interest in this particular problem.

