## Supplementary Materials for "A fossilized birth-death model for fossil records lacking sampled ancestors"

### Verifying the analytical FBDT likelihood

We verified that the analytical FBDT likelihood derived in the Appendix is numerically equivalent to the ODE-based implementation used by `MiSSE()`. The code below evaluates both likelihoods using the same simulated tree, fossil data, and parameter values, and demonstrates that they return identical log-likelihoods (up to numerical precision).

First, load the required packages (`hisse` and `diversitree`) and simulate a simple birth–death tree with fossil sampling:

```
set.seed(8)
phy <- TreeSim::sim.bd.taxa(n = 100, numbsim = 1, lambda = 0.3,
    mu = 0.2)[[1]]
f <- hisse:::GetFossils(phy, psi = 0.05)
pp <- hisse:::ProcessSimSample(phy, f)
```

Next, organize the simulated data into the format required by the internal `hisse` functions:

```
dat.tab <- hisse:::OrganizeDataMiSSE(phy = pp$phy, f = 1, hidden.states = 2,
    includes.fossils = TRUE)
edge_details <- hisse:::GetEdgeDetails(phy = pp$phy, intervening.intervals = strat.cache$intervening.in
fossil.taxa <- edge_details$tipward_node[which(edge_details$type ==
    "extinct_tip")]
# This is for starting values:
fossil.ages <- dat.tab$TipwardAge[which(dat.tab$DesNode %in%
    fossil.taxa)]
# Remove k fossils so that the remaining tree contains only
# extant taxa and observed fossil tips. Split times are
# then calculated from this reduced tree.
k.sample.tip.no <- grep("Ksamp*", x = phy$tip.label)
phy.no.k <- drop.tip(pp$phy, k.sample.tip.no)
split.times <- paleotree:::dateNodes(phy.no.k, rootAge = max(node.depth.edgelength(phy.no.k)))[-c(1:Nti
# n = number of extant taxa
n <- Ntip(phy.no.k) - length(fossil.taxa)
# m = number of observed fossil tips
m <- length(fossil.taxa)
x_times <- split.times
y_times <- fossil.ages
k <- 0
```

We next evaluate the analytical FBDT log-likelihood using the generating parameter values from the simulation:

```
rho = 1
lambda <- 0.3
mu <- 0.2
psi <- 0.05
# Note the removal of k from this equation and the use
# p_one_censored().
logLikLogSpace <- (((n + m - 2) * log(lambda)) + (m * log(psi))) -
    log(1 - hisse:::p_0(max(x_times), lambda, mu, psi = 0, rho,
        log = FALSE)) * 2 + hisse:::p_one_censored(max(x_times),
    lambda, mu, psi, rho) + sum(hisse:::p_one_censored(x_times,
    lambda, mu, psi, rho)) + (sum(hisse:::p_0(y_times, lambda,
    mu, psi, rho)) - sum(hisse:::p_one_censored(y_times, lambda,
    mu, psi, rho)))
```

We then evaluate the likelihood using the ODE-based routine employed by `MiSSE()`:

```
phy <- pp$phy
nb.tip <- Ntip(phy)
nb.node <- phy$Nnode
dat.tab <- hisse:::OrganizeDataMiSSE(phy = phy, f = 1, hidden.states = 2,
    includes.fossils = TRUE)
model.vec <- c(lambda + mu, mu/lambda, lambda + mu, mu/lambda,
    rep(0, 48), 0.01, psi)
cache = hisse:::ParametersToPassMiSSE(model.vec = model.vec,
    hidden.states = 1, fixed.eps = NULL, nb.tip = nb.tip, nb.node = nb.node,
    psi.type = "m_only", bad.likelihood = exp(-300), ode.eps = 0)
gen <- hisse:::FindGenerations(phy)
k.samples <- NULL
MiSSE.logL <- hisse:::DownPassMisse(dat.tab = dat.tab, cache = cache,
    gen = gen, condition.on.survival = TRUE, root.type = "madfitz",
    root.p = NULL, fossil.taxa = fossil.taxa, node = NULL, fix.type = NULL)
```

Finally, we compare the two log-likelihoods:

```
analytical <- round(logLikLogSpace, 3)
ode <- round(MiSSE.logL, 3)
analytical
```

```
## [1] -473.065
```

```
ode
```

```
## [1] -473.065
```

```
analytical == ode
```

```
## [1] TRUE
```

The comparison returns `TRUE`, confirming that the analytical and ODE-based implementations produce identical log-likelihoods (up to numerical precision).
